# Generative Adversarial Implicit Successor Representation

**DOI:** 10.1101/2024.03.03.581078

**Authors:** Whan-Hyuk Choi, Sukyeon Ha, Hansol Choi

## Abstract

We propose a novel method to implicitly encode Successor Representations (SRs) using a Generative Adversarial Network (GAN). SRs are a method to encode states of the environment in terms of their predictive relationships with other states, which can be used to predict long-term future rewards. In standard explicit methods, the value of SR is found from an explicit map between future states after an action or to find an approximate function. Instead, our method encodes SR implicitly using a GAN. The distribution of samples generated by the GAN system approximates the successor representation. We also suggest an action decision procedure for the implicit encoding of SR. The system makes the decision using an analysis-by-synthesis procedure that it attempts to synthesize a sample that can explain the action decision constraints of the current and target states. Our system is different from the classical SR in several points. It can sample actual samples reflecting SR distribution, which is not easy for explicit models. It can also get around the issue of explicitly representing probabilities or successor representation values and doing math over them. We tested our system in a toy environment, where the agent could learn the implicit successor representation successfully and use it for action decisions.

## 1 INTRODUCTION

Successor Representation (SR) is a recently developed method in reinforcement learning studies. SR is to capture the expected future state occupancy of an agent, following its state and action at that time. As a result, it encodes the long-term effects of actions in the environment. (Dayan, 1993; Kulkarni et al., 2016; Zhang et al., 2017) It is particularly useful in complex tasks where the immediate effects of actions are not always indicative of their long-term consequences. SRs, especially with deep learning estimation method called successor features(Barreto et al., 2017), have been applied to a variety of different problems and domains within RL. It demonstrates their versatility and effectiveness. SR is not only important in machine learning but also recently suggested as a computational model to explain the brain’s memory models. The hippocampus is suggested to use SR to encode the state spaces (Nyberg et al., 2022; Stachenfeld et al., 2017). SR could also be used to explain the behavior of humans in reward-changing environments (Russek et al., 2017; Momennejad et al., 2017).

Recent advances in generative adversarial network (GAN) provided a method to train implicit generative models (Goodfellow et al., 2014). GAN is an unsupervised method to train a model that can generate samples resembling target data (Radford et al., 2015). In an optimal learning condition, GAN can generate samples whose distribution is identical to the training dataset and represent it implicitly. It is distinguishable from explicit generative models, which have specification of explicit functions and parameters to specify (Gershman and Beck, 2017; Mohamed and Lakshminarayanan, 2017). GAN is a useful way to train implicit generative models because it is said to automatically find and learn patterns in a lot of different types of input data, such as pictures, videos, and sounds (see (Creswell et al., 2018) for more information).

We propose an alternative encoding for SR, named Generative Adversarial Implicit Successor Representation (GAISR). GAISR produces samples whose distribution reflects the SR values; more samples are generated for larger SR values. We also suggest an action-decision procedure using GAISR. As explained above, an explicit SR system uses the SR values to compute the value of actions. In our GAISR, the action decision is done by sample generation and sample selection procedures. It is similar to the in-painting method by Yeh et al. (Yeh et al., 2016). GAISR generates random samples using a trained GAN as a result, the generated samples reflect the distribution of the training data. Filtering the samples for the specified constraints comes next. We will explain the GAISR architecture, and a toy-model demonstration will be followed.

### 1.1 Relationship with Existing Works

Kulkarni et al. (Kulkarni et al., 2016) used a deep network for encoding successor representation. Their purpose was to compute the SR of a states represented in high-dimensional spaces. Deep networks reduce state space to smaller feature space. The method, successor feature representation, has been widely used for machine learning studies.(Zhang et al., 2017) Milledge and Buckley recently suggested active inference successor representation (Millidge and Buckley, 2023), in which they used SR to represent future space to compute the variational free energy. Both method is different from our study that they encode the SR value explicitly.

## 2 MATERIALS AND METHODS

### 2.1 GAISR and action inference

#### 2.1.1 Training GAISR

SR is the expected frequency of visits to a state *S*′ along a sequence of states starting with a state and action (*S, A*), with policy *π*. The basic SR is defined by discrete states and action spaces. Discrete states are expressed with capital letters such as *S, A*.

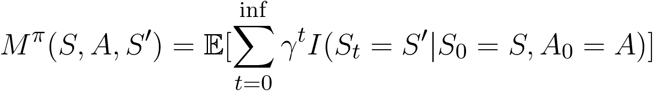

The parameter *γ* discounts the number of visits by time between the states. Our method, GAISR is to encode SR implicitly. A generator function *G*(*z*) takes noise *z* from a distribution *Z* and projects it to (*s, a, s*′) = *G*(*z*). *s, a, s*′ are current state, action and the future states defined by continuous values. GAISR encodes the SR by letting the frequency of its generated samples reflect *M* (*S, A, S*′).

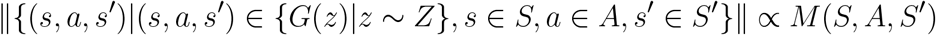

∥ · ∥ indicate the number of samples in the set. *s* ∈ *S* indicates that the continuous state and action *s, a, s*′ are instance of the discrete state and action *S, A* and *S*′ correspondingly.

GAISR is composed of a working memory (WM) and a GAN encoder (Fig. 1). WM stores the history of the states and actions (*s, a*) from the agents’ history of world visits. WM generates the training data of (*s, a, s*′) to the following GAN memory. To reflect the discount factor of SR, the working memory decays by time with a discount factor *γ. P* ((*s, a, s*′) ∈ *D*_*train*_) = *γ*^*τ*^, where *τ* is the time between *s* and *s*′. *D*_*train*_ is the training set data for GAN. The memory drops out samples by the chance of 1 *− γ* for each iteration to discount the visitation information following the definition of standard SR.

**Figure 1.**
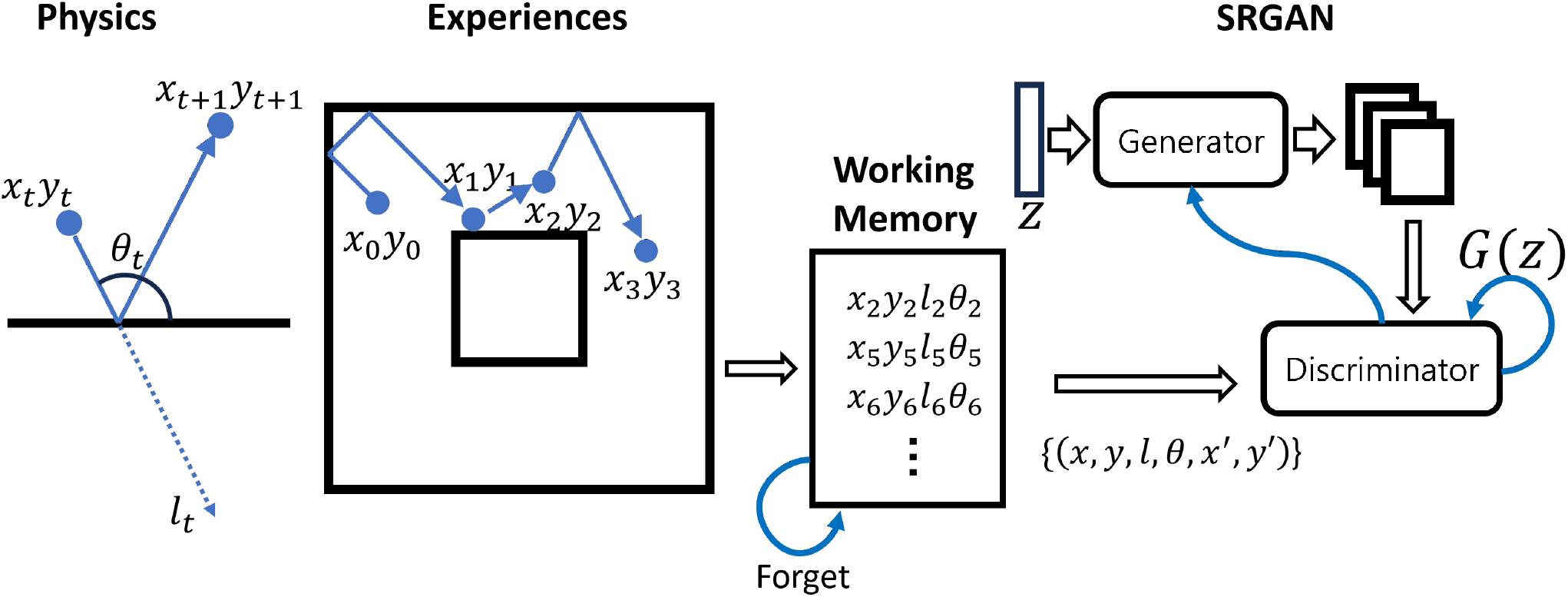
The general structure of SRGAN system. The experiences from environments are stored in working memory, which memorizes previous state and action pairs. The working memory forgets its entry by the chance of *γ* for each step. The distribution of samples of previous state, action, and current state pairs generated with WM is correlated with the SR of the environment. GAN captures the distribution of the samples.

GAN learned to implicitly encode the distribution of the training data from WM. It can be considered a long-term memory of our GAISR. GAN is composed of two networks. (Goodfellow et al., 2014; **?**; Yeh et al., 2016). The generator *G* receives noise *z* from a distribution *z Z* and maps *G*(*z*) to the sample space: (*s, a, s*′). Discriminator *D* receives input from *G* and *D*_*train*_.

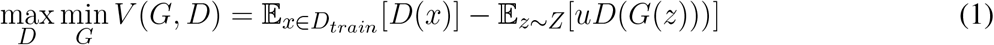

#### 2.1.2 Action decision as sample generation and filter

We suggest PAI algorithm using a trained GAISR. In a standard SR model, SR system will compute the Q-value of each state and action with simple computation. 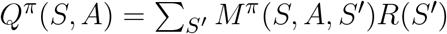 Where *R*(*S*′) is the reward of state *S*′. Instead, our method infers the action to achieve 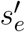 in the future of *s*_*e*_ by analysis-by-synthesis manner. It generates samples {(*s, a, s*′) = *G*(*z*)|*z ∼ Z}* using trained generator *G*.

The *z* with minimum distance between its sample and given evidences of current state and target state is chosen as the optimal. 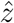. The action *a* from. 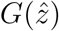 is chosen as the optimal action.

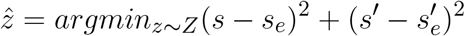

The GAISR agent uses the action *â* is the action from. 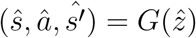

### 2.2 Simulation

Fig. 1 shows our toy environment and the architecture of GAISR framework. The objective of the GAISR agent in the toy environment is to move toward a target state, which requires multiple steps of actions. The toy environment provides experience. For each time *t*, the agent is at a state, *s*_*t*_ = (*x*_*t*_, *y*_*t*_) defined in a Cartesian environment. The agent shoots to *s*_*t*+1_ = (*x*_*t*_ + 1, *y*_*t*_ + 1). The shooting action happens with the direction and the shooting length of *a*_*t*_ = (*θ*_*t*_, *l*_*t*_). The environment has walls (black, thick lines). When the agent collapses into a wall, it reflects back in a perfect elastic collision. The agents move around in our doughnut-shaped toy environment. One iteration of movement is composed of multiple shooting events. Each iteration begins from a uniformly random position in the doughnut.

The agent initially used a random null policy *π*^0^ to explore the environment and use the experience to build an SR map of GAISR^0^. The agent with *π*_1_ uses the GAISR^0^ to generate actions.

## 3 RESULTS

### 3.1 Training GAISR

We let our system travel around the doughnut environment with a random policy *π*^0^, with random actions for 10000 iterations, each composed of 10 shoot sequences. Each iteration starts from a random position in the environment. Each shoot starts from the previous end position. Each iteration starts from a random position in the environment. GAISR was trained using the history of the data.

After training, we first check the structure of the generated samples. Samples were generated from the trained generator *{G*(*z*)|*z ∼ Z*}. After training, the samples reflect the structure of the donut-shaped space (Fig. 2 A). On the other hand, the naive and early stage GAISR (Supplementary Fig. 1) does not have the structure. The trained GAISR became closer to the training data from the working memory (Fig. 2 A).

**Figure 2.**
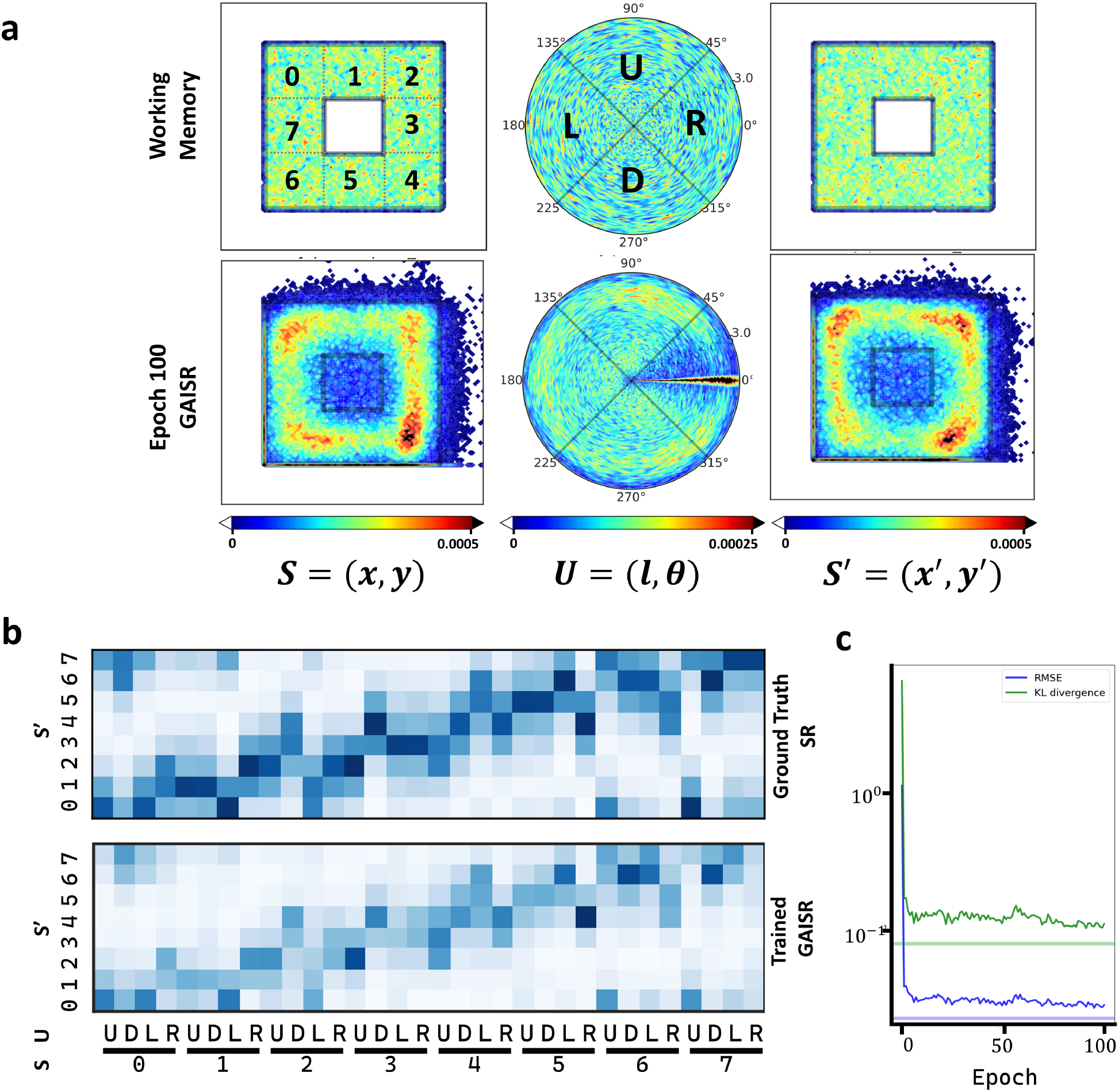
**a** The sample distribution(n=100,000) from the trained GAISR and the training data. The training data is from the working memory. The samples generated by Trained GAISR reflects the environmental structures. The current and future states avoids the empty center space of the doughnut shaped environment and the actions are spread over the whole direction and length. **b** Trained GAISR generates samples reflecting the successor representation of the environment. The distribution of samples from GAISR is similar to the ground truch SR estimated by Monte Carlo method. The state and action space was divided into discrete states and actions for visualization and comparison purpose. **c** Training GAISR let the distribution of SR close to the ground truth SR. The horizontal lines shows the difference between the ground truth SR and the distribution of training data from working memory. They are the limit of training.

Our GAN system implicitly encoded SR (Fig. 2). The trained GAISR generated samples whose distribution is close to the ground truth SR, estimated by Monte-Carlo method (Fig. 2 B). The SR distribution was driven by the training data from the working memory (Supplementary Fig. 1). The samples from an untrained network or the very early stage of training (Supplementary Fig. 2) were different from ground truth SR(Supplementary Fig. 3). The samples from *GAISR* shows the structure of the environment and policy. For example, the SR of right action from *state*0 is biased toward *state*2, as expected from the random policy of the agent. We use the distribution of samples from GAISR to estimate the SR and used the estimated SR to compute Q-values. The Q-values became structuraly close to the ground truth Q-values (Supplementary Fig 3.)

**Figure 3.**
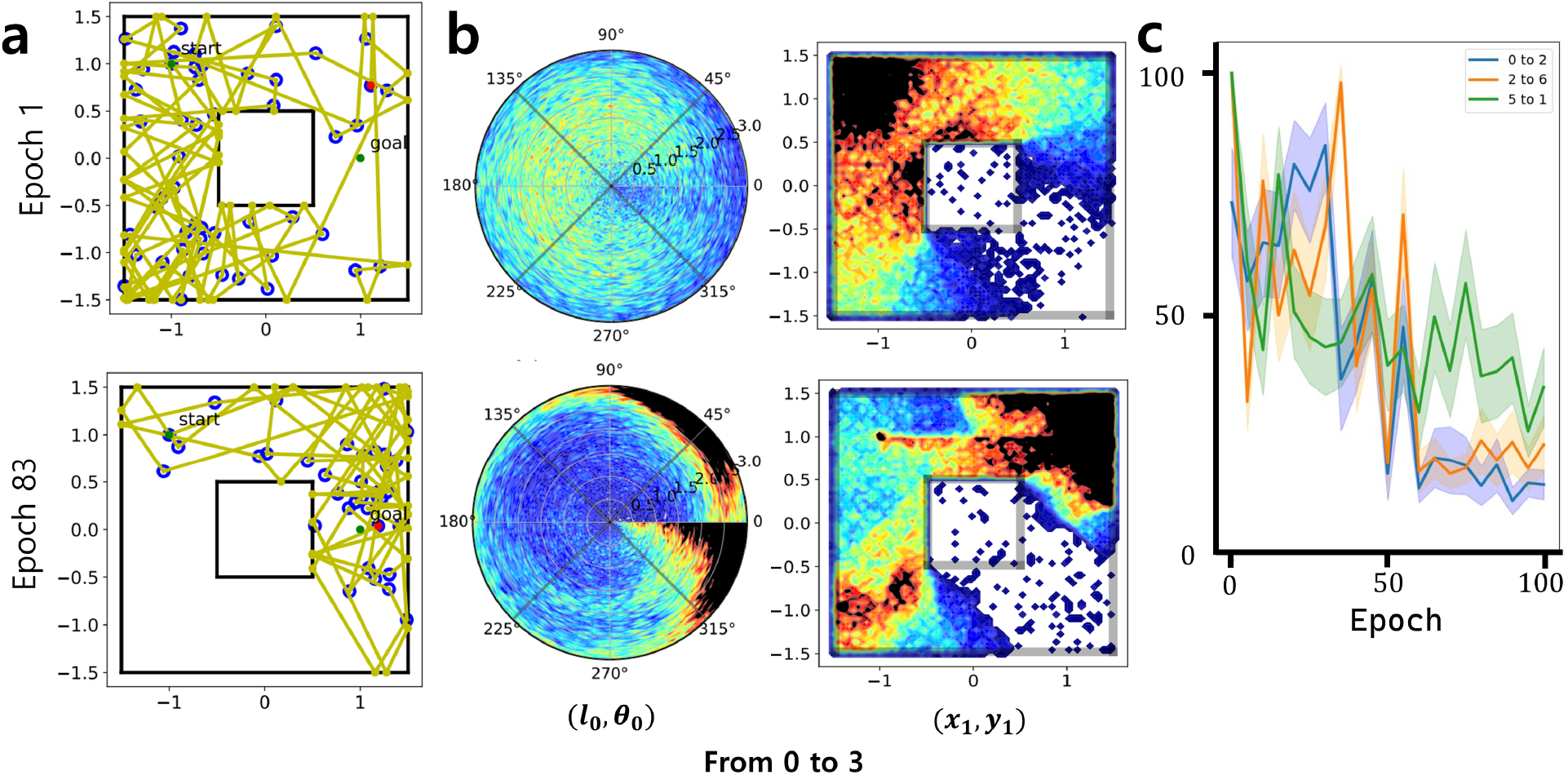
Inference of Action with SRGAN. **a** Representative movement of an agent following the action inference, starting from the center of state 0 and reaching the center of state 3. The path of agent with median number of actions at the early and the later stage of training is shown. **b** Distribution of actions and their resulting states(n=100,000) at the first step of inference, from state 0 to 3. The trained model shows actions biased toward direction of right and length of 3, which is preferable to reach the goal state. **c** The average number of steps needed to reach the target area(circle, r=0.2), with 95 percent confidence interval. Three different combination of start and end positions are tested, all of which consistently showed improvement through training. If a trial didn’t reach the target within 50 steps, the trial is considered to take 100 steps to reach.

### 3.2 Inference of actions using SRGAN

GAISR infers the actions under an analysis-by-synthesis method. When current state *s*_*o*_ = (*x*_*o*_, *y*_*o*_) and the target state .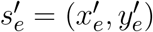 are given, the GAISR tries to generate sample, whose current and future states match the given current and target state. This is done by generating samples and selecting the sample with the minimum distance to the given evidences. The agent acts with the action of the optimal sample. Fig. 3A. shows a representative result from an untrained and trained GAISR system. The system continues to move until it reaches either the goal state or the step limit of 50. The distribution of actions at the initial state and the distribution of states at the next step show the difference between early-stage and late-stage training GAISR (Fig. 3 B). The action is widely spread over the whole direction and length in the early stage training, while it gets densely concentrated to the most effective length(3) and direction(right) to reach the goal state in the later stage training. The totally naiive initial GAISR can generate only limited action repertoire (Supplementary Fig. 4) Trained GAISR shows less number of actions to achieve the target state, compared to untrained GAISR (Fig. 3C).

## 4 DISCUSSION

In a previous study (Lenninger et al., 2023), we suggested a method to use GAN to implicitly encode the density of the state-action-state sequence of a multi-joint arm system. We used the generative model to solve the forward and inverse problems of motor control. The result was limited by time, as the system could encode only a single step. This study extends the previous work to infer actions toward a not-adjacent future target. We added a working memory system to store the history of states with time decay for this purpose. As a result, our implicit generative model became possible to generate samples, with their density encoding the SR map of our environment. We also suggested the inference method in GAISR. Explicit methods such as Bayesian statistics need to integrate prior and likelihood to compute posterior. Instead, our method is defined as procedures of generating samples.

SRs are not only used for machine learning but also suggested as the functional model for brains. Important questions are: how the brains encode and learn SR values, and use them for action. In the previous studies, the SR was modeled explicitly. When you have a state and a future state pair, you can find a SR value encoded in a map or a function. Learning is to update the map or the parameters of the approximation function. The system use the SR values numerically to compute values of states or actions for decision-making (Gershman, 2019; Mohamed and Lakshminarayanan, 2017). Numerical encoding of values can well explain the experimental evidence, but, the question of encoding and computing values in neural circuits remains (Sohn and Narain, 2021). The implicit generative models may provide an operational algorithm for the numerical idea that, they can directly encode the samples, and the inference can be made by the sample generation procedure of analysis-by-synthesis.

Adversarial dual learning is not a new idea in modeling brain functions.For example, a dual learning model in RL, actor-critic model has been suggested to explain the function of DAergic systems (Houk et al., 1995; Dunovan and Verstynen, 2016). The DA signal have been considered to encode reward-prediction error (Schultz et al., 1997) but recent evidences shows that it can encode a general state prediction error signal (Nakahara et al., 2004; Gardner et al., 2018; Noudoost and Moore, 2011; Suarez et al., 2019; Takahashi et al., 2017; Puig et al., 2014; Gershman and Schoenbaum, 2017). DAergic output to thalamus is considered a perception or action decision problems (Ding and Gold, 2013; Lak et al., 2017; Wei et al., 2015; Babayan et al., 2018; Keuken et al., 2015). DA signal teaches both the cortex and basal ganglia systems. (Joel et al., 2002) Those are necessary properties of the discriminator in GAN. Interestingly, Pfau et al. (Pfau and Vinyals, 2016) also suggested the architectural similarity between GAN and actor-critic models.Gershman recently suggested adversarial generative brain hypothesis. The brains are suggested to use the generative-adversarial for training the implicit generative model of the brains. (Gershman, 2019) We suggest an adversrarial learning framework, focusing on the sample generation procedure.

We expect that this implicit SR model can be used in the practical decision-making problems, which needs actual sample generation. Especially, GAN has been shown to be successful to train a generative model for complex conditions. Our GAISR might also be extendable for problems needing complex generative models. Also, we expect that our model be used to explain the learning and inference procedures of SR brains’ models and to provide the underlying procedures of the statistical models. Our model can be considered as a procedural implementation for planning-as-inference framework (Botvinick and Toussaint, 2012; Solway and Botvinick, 2012) suggested in computational neurosciences. Their objective is to infer the probability of current action *a* with the current state *s* and the goal state *s*′ as the evidence of inference. It uses the generative model of state and action sequences for inference. The process of GAISR can be concerned as an analysis-by-synthesis (Yuille and Kersten, 2006) method. An agent with a generative model can use it for perception by trying to generate samples that can best explain the given observation. Our method is to generate sample that can explain the current and target states under an SR generative model.

## AUTHOR CONTRIBUTIONS

WC conceptualized, analysed, acquired funding, investigated the results, and wrote the manuscript. SH developed the software, analysed the data, wrote the manuscript, investigated the results, and HC conceptualized, designed, and supervised the project and wrote the manuscript.

## REFERENCES

Babayan, B. M., Uchida, N., and Gershman, S. J. (2018). Belief state representation in the dopamine system. Nature Communications 9, 1891. doi:10.1038/s41467-018-04397-0

Barreto, A., Dabney, W., Munos, R., Hunt, J. J., Schaul, T., van Hasselt, H. P., et al. (2017). Successor features for transfer in reinforcement learning. In Advances in Neural Information Processing Systems (Curran Associates, Inc.), vol. 30

Botvinick, M. and Toussaint, M. (2012). Planning as inference. Trends in Cognitive Sciences 16, 485–488. doi:10.1016/j.tics.2012.08.006.00020

Creswell, A., White, T., Dumoulin, V., Arulkumaran, K., Sengupta, B., and Bharath, A. A. (2018). Generative adversarial networks: An overview. IEEE Signal Processing Magazine 35, 53–65. doi:10.1109/MSP.2017.2765202

Dayan, P. (1993). Improving generalization for temporal difference learning: The successor representation. Neural Computation 5, 613–624. doi:10.1162/neco.1993.5.4.613

Ding, L. and Gold, J. (2013). The basal ganglia’s contributions to perceptual decision making. Neuron 79, 640–649. doi:10.1016/j.neuron.2013.07.042

Dunovan, K. and Verstynen, T. (2016). Believer-skeptic meets actor-critic: Rethinking the role of basal ganglia pathways during decision-making and reinforcement learning. Frontiers in Neuroscience 10. doi:10.3389/fnins.2016.00106

Gardner, M. P. H., Schoenbaum, G., and Gershman, S. J. (2018). Rethinking dopamine as generalized prediction error. Proceedings of the Royal Society B: Biological Sciences 285, 20181645. doi:10.1098/rspb.2018.1645

Gershman, S. J. (2019). The generative adversarial brain. Frontiers in Artificial Intelligence 2. doi:10.3389/frai.2019.00018

Gershman, S. J. and Beck, J. M. (2017). Complex Probabilistic Inference (John Wiley and Sons Ltd). 453–466. doi:10.1002/9781119159193.ch33

Gershman, S. J. and Schoenbaum, G. (2017). enRethinking dopamine prediction errors. bioRxiv, 239731doi:10.1101/239731

Goodfellow, I., Pouget-Abadie, J., Mirza, M., Xu, B., Warde-Farley, D., Ozair, S., et al. (2014). Generative Adversarial Nets (Curran Associates, Inc.). 2672–2680

Houk, J. C., Adams, J. L., and Barto, A. G. (1995). A model of how the basal ganglia generate and use neural signals that predict reinforcement (The MIT Press). Computational neuroscience. 249–270

Joel, D., Niv, Y., and Ruppin, E. (2002). Actor–critic models of the basal ganglia: new anatomical and computational perspectives. Neural Networks 15, 535–547. doi:10.1016/S0893-6080(02)00047-3

Keuken, M. C., Maanen, L. V., Bogacz, R., Schäfer, A., Neumann, J., Turner, R., et al. (2015). The subthalamic nucleus during decision-making with multiple alternatives. Human Brain Mapping 36, 4041–4052. doi:10.1002/hbm.22896

Kulkarni, T. D., Saeedi, A., Gautam, S., and Gershman, S. J. (2016). Deep successor reinforcement learning doi:10.48550/arXiv.1606.02396. 1606.02396 [cs, stat]

Lak, A., Nomoto, K., Keramati, M., Sakagami, M., and Kepecs, A. (2017). Midbrain dopamine neurons signal belief in choice accuracy during a perceptual decision. Current Biology 27, 821–832. doi:10.1016/j.cub.2017.02.026

Lenninger, M., Choi, W.-H., and Choi, H. (2023). enPredictive motor control based on a generative adversarial network, 2023.01.17.524156doi:10.1101/2023.01.17.524156

Millidge, B. and Buckley, C. L. (2023). enActive Inference Successor Representations (Cham: Springer Nature Switzerland), vol. 1721 of Communications in Computer and Information Science. 151–161. doi:10.1007/978-3-031-28719-011

Mohamed, S. and Lakshminarayanan, B. (2017). Learning in implicit generative models. 1610.03483 [cs, stat] ArXiv: 1610.03483

Momennejad, I., Russek, E. M., Cheong, J. H., Botvinick, M. M., Daw, N. D., and Gershman, S. J. (2017). enThe successor representation in human reinforcement learning. Nature Human Behaviour 1, 680–692. doi:10.1038/s41562-017-0180-8

Nakahara, H., Itoh, H., Kawagoe, R., Takikawa, Y., and Hikosaka, O. (2004). Dopamine neurons can represent context-dependent prediction error. Neuron 41, 269–280. doi:10.1016/S0896-6273(03)00869-9

Noudoost, B. and Moore, T. (2011). Control of visual cortical signals by prefrontal dopamine. Nature 474, 372–375. doi:10.1038/nature09995

Nyberg, N., Duvelle,, Barry, C., and Spiers, H. J. (2022). Spatial goal coding in the hippocampal formation. Neuron 110, 394–422. doi:10.1016/j.neuron.2021.12.012

Pfau, D. and Vinyals, O. (2016). Connecting generative adversarial networks and actor-critic methods. arXiv:1610.01945 [cs, stat] ArXiv: 1610.01945

Puig, M. V., Rose, J., Schmidt, R., and Freund, N. (2014). Dopamine modulation of learning and memory in the prefrontal cortex: insights from studies in primates, rodents, and birds. Frontiers in Neural Circuits 8. doi:10.3389/fncir.2014.00093

Radford, A., Metz, L., and Chintala, S. (2015). Unsupervised representation learning with deep convolutional generative adversarial networks. 1511.06434 [cs] ArXiv: 1511.06434

Russek, E. M., Momennejad, I., Botvinick, M. M., Gershman, S. J., and Daw, N. D. (2017). enPredictive representations can link model-based reinforcement learning to model-free mechanisms. PLOS Computational Biology 13, e1005768. doi:10.1371/journal.pcbi.1005768

Schultz, W., Dayan, P., and Montague, P. R. (1997). A neural substrate of prediction and reward. Science (New York, N.Y.) 275, 1593–9

Sohn, H. and Narain, D. (2021). Neural implementations of bayesian inference. Current Opinion in Neurobiology 70, 121–129. doi:10.1016/j.conb.2021.09.008

Solway, A. and Botvinick, M. M. (2012). Goal-directed decision making as probabilistic inference: A computational framework and potential neural correlates. Psychological Review 119, 120–154. doi:10.1037/a0026435

Stachenfeld, K. L., Botvinick, M. M., and Gershman, S. J. (2017). enThe hippocampus as a predictive map. Nature Neuroscience 20, 1643–1653. doi:10.1038/nn.4650

Suarez, J. A., Howard, J. D., Schoenbaum, G., and Kahnt, T. (2019). Sensory prediction errors in the human midbrain signal identity violations independent of perceptual distance. eLife 8, e43962. doi:10.7554/eLife.43962

Takahashi, Y. K., Batchelor, H. M., Liu, B., Khanna, A., Morales, M., and Schoenbaum, G. (2017). Dopamine neurons respond to errors in the prediction of sensory features of expected rewards. Neuron 95, 1395–1405.e3. doi:10.1016/j.neuron.2017.08.025

Wei, W., Rubin, J. E., and Wang, X.-J. (2015). Role of the indirect pathway of the basal ganglia in perceptual decision making. Journal of Neuroscience 35, 4052–4064. doi:10.1523/JNEUROSCI.3611-14.2015

Yeh, R., Chen, C., Lim, T. Y., Hasegawa-Johnson, M., and Do, M. N. (2016). Semantic image inpainting with perceptual and contextual losses. 1607.07539 [cs] arXiv: 1607.07539

Yuille, A. and Kersten, D. (2006). Vision as bayesian inference: analysis by synthesis? Trends in Cognitive Sciences 10, 301–308. doi:10.1016/j.tics.2006.05.002

Zhang, J., Springenberg, J. T., Boedecker, J., and Burgard, W. (2017). Deep reinforcement learning with successor features for navigation across similar environments. In 2017 IEEE/RSJ International Conference on Intelligent Robots and Systems (IROS). 2371–2378. doi:10.1109/IROS.2017.8206049

