## Supplementary materials for "Generative Adversarial Implicit Successor Representation"

#### Supplementary Material

### 1 SUPPLEMENTARY TABLES AND FIGURES

#### 1.1 Figures

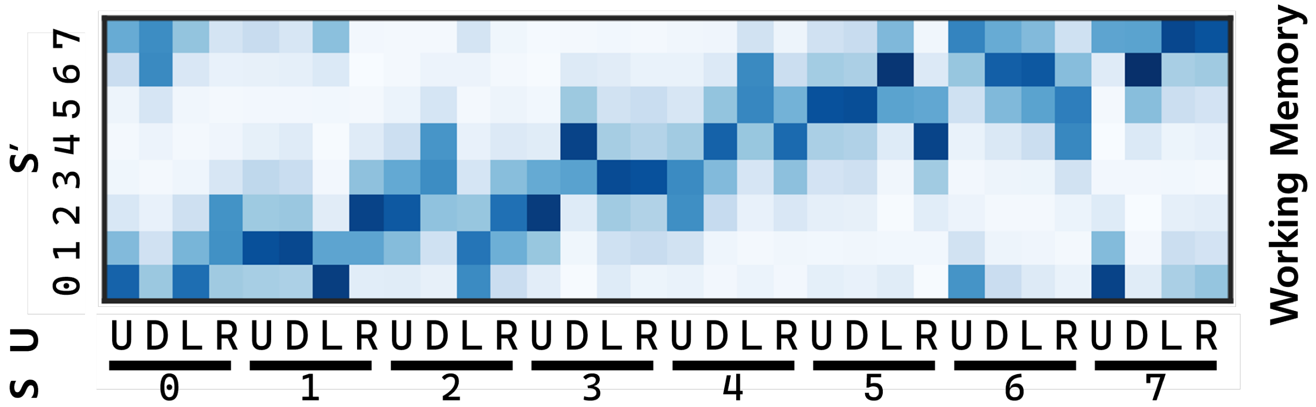


**Figure S1.** The distribution of samples from the working memory. The samples are used for training GAISR. The structure of this distribution reflect the SR of the agent. The working memory tracks current state, current action, and the future state from the state and action. The data from working memory reflects the temporal decay of memory.

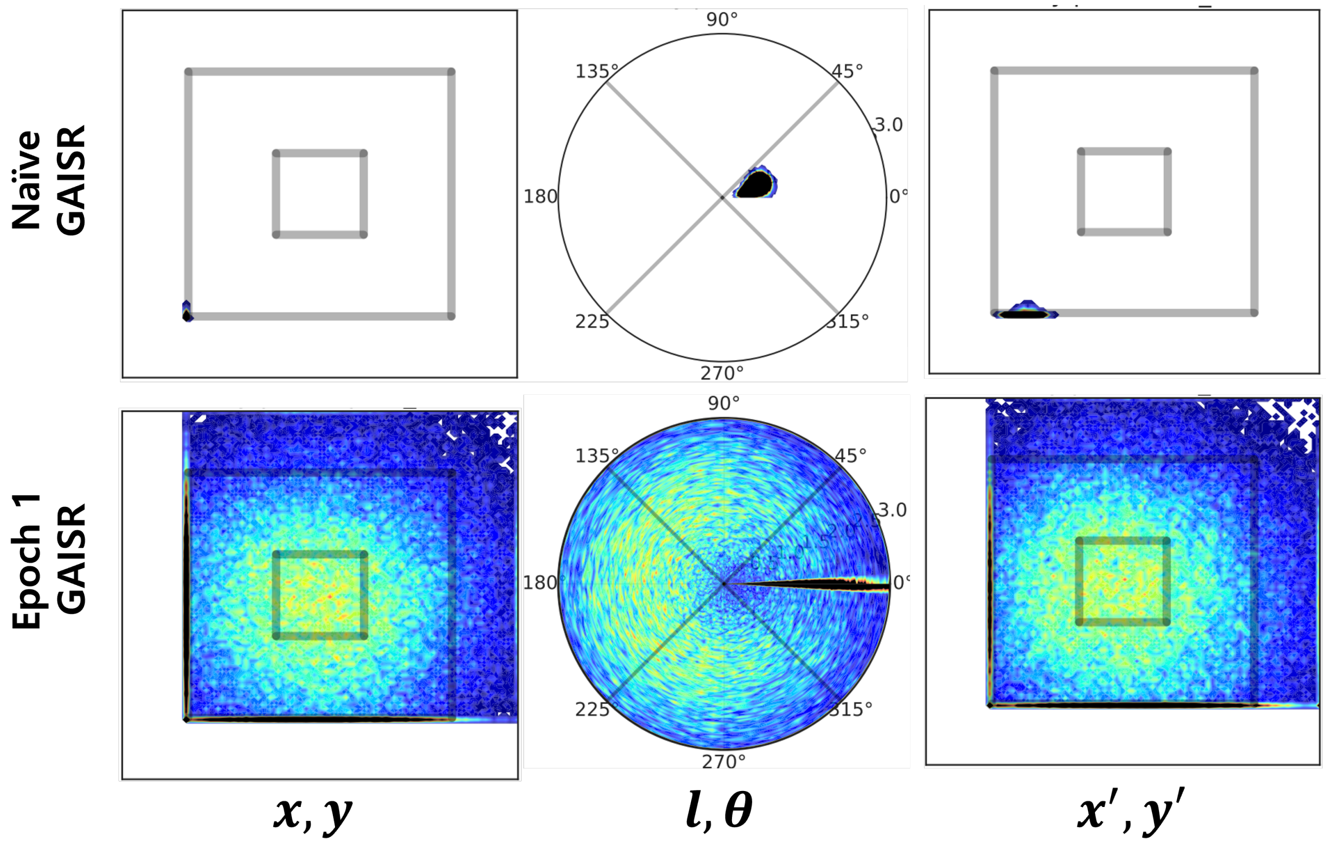


**Figure S2.** The distribution of samples from the naive (training epoch 0) and the early stage training (epoch 1) GAISR. They do not encode the SR of the environment. The samples from the naive model are highly biased to a single sample. The samples from the early stage model shows lots of samples in the center of doughnut, which is not allowed for the agent to stay.

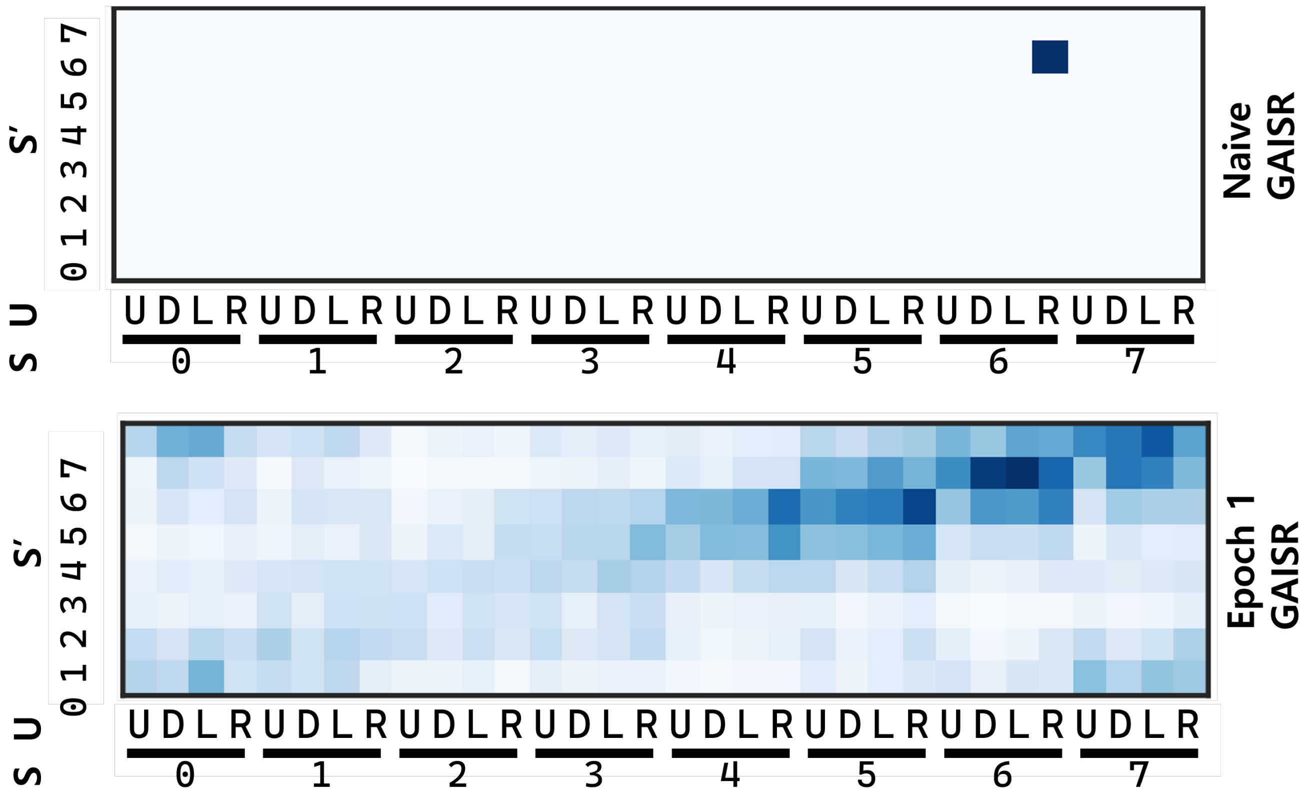


**Figure S3.** The distribution of samples from the naive (epoch 0) and early stage training (epoch 1) GAISR. They do not reflect the ground-truth SR.

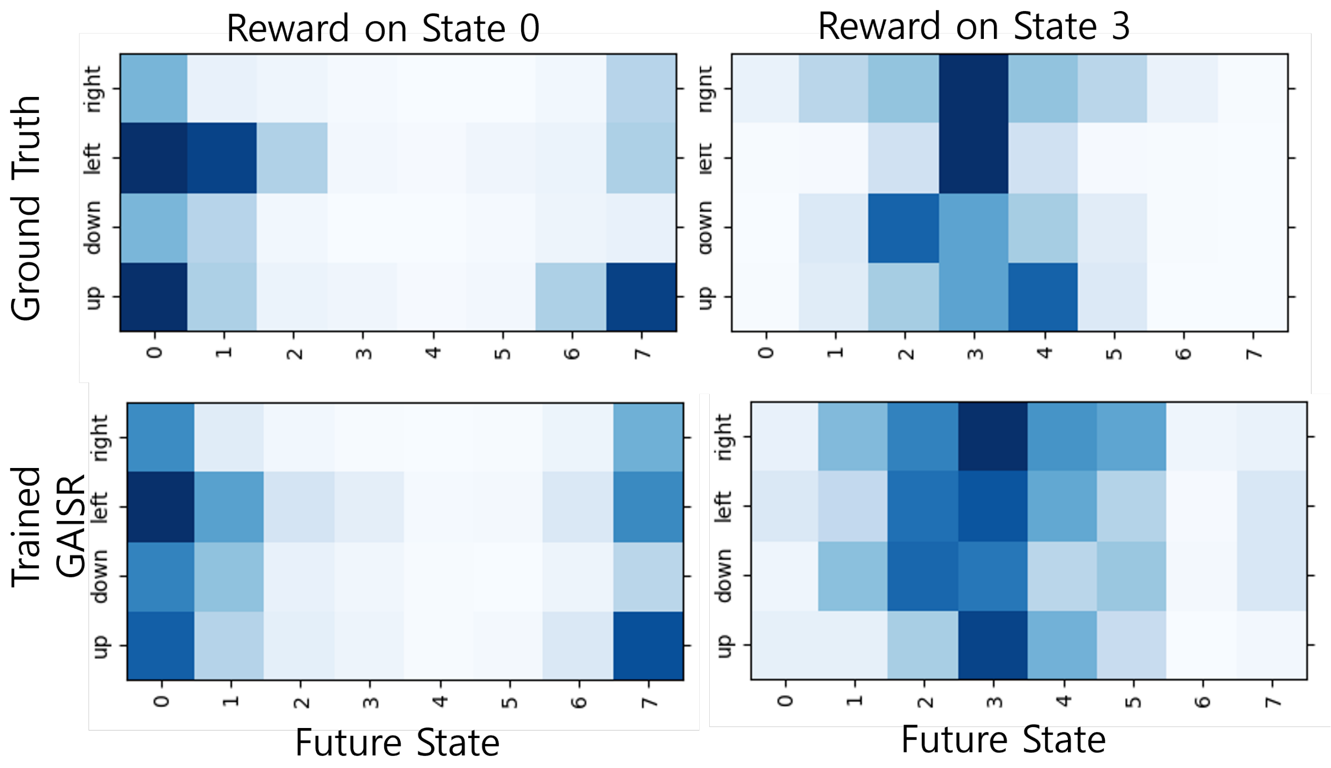


**Figure S4.** Q-values estimated from the distribution of samples generated by trained GAISR. Their sturcturs are comparable with the ground truth Q-values. For each of the figure, the Q-value $Q(S, A)$ is shown, in case of reward is located at state 0 or state 3. You can calculate the Q-value using the formula $Q(S, A) = sum'_S M(S, A, S')R(S')$. The ground truth Q-value is computed with SR from a Monte-Carlo simulation of the agent's behavior. The $M$ of trained GAISR is from the Monte-Carlo sampling on GAISR.

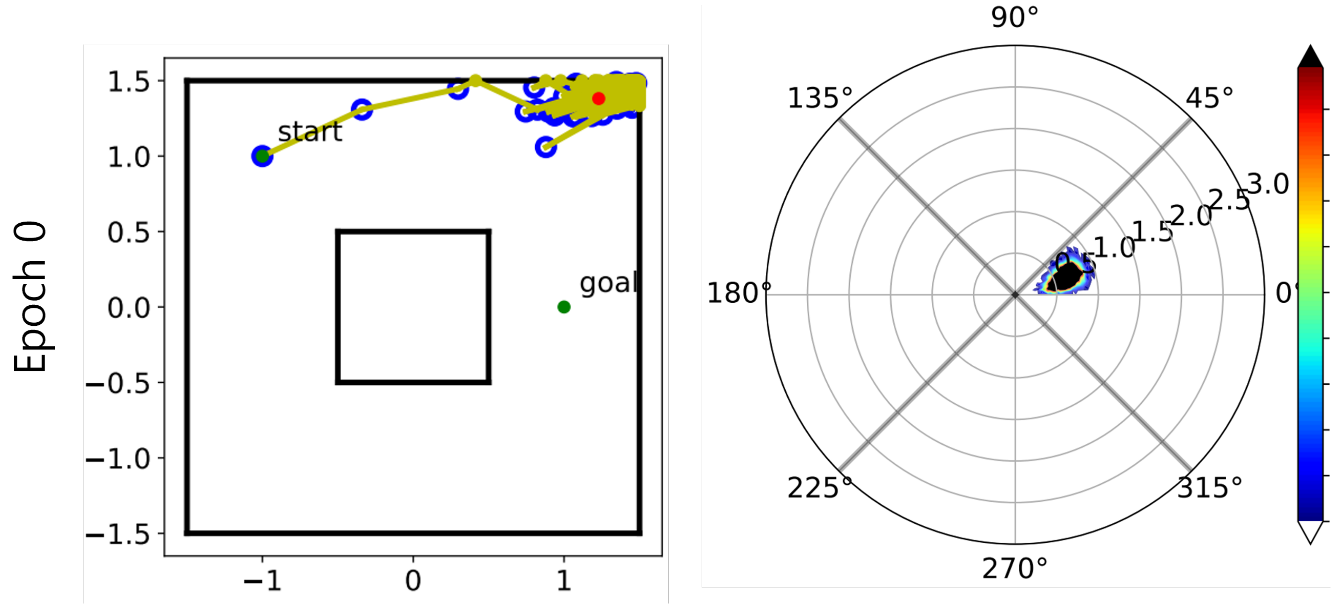


**Figure S5.** The sequence of actions using a naive GAISR. The actions are highly biased in one direction, regardless of the current state. The agent could not reach the target.
